# A new clade of pararetroviruses distantly related to hepadnaviruses and nackednaviruses

**DOI:** 10.1101/2024.08.02.606351

**Authors:** Jaime Buigues, Adrià Viñals, Raquel Martínez-Recio, Juan S. Monrós, José M. Cuevas, Rafael Sanjuán

## Abstract

Group VII of the Baltimore classification comprises reverse-transcribing, non-integrated DNA viruses, also known as pararetroviruses. These include the hepadnaviruses, a family of small enveloped DNA viruses that infect vertebrates, but also a sister family of non-enveloped fish viruses, the nackednaviruses. Here we describe the complete sequence of a new pararetrovirus found in the feces of an insectivorous bat. This virus encodes a core protein and a reverse transcriptase but no envelope protein. A database search identified a viral sequence from a permafrost sample as its closest relative. The two viruses form a cluster that occupies a basal phylogenetic position relative to hepadnaviruses and nackednaviruses, with an estimated divergence time of 500 million years. These findings may lead to the definition of a new viral family and support the hypothesis that ancestral animal pararetroviruses were non-enveloped.

## Main text

Two major families of pararetroviruses have been defined by the International Committee for the Taxonomy of Viruses: *Caulimoviridae* (1) and *Hepadnaviridae* (2). While caulimoviruses infect plants, hepadnaviruses infect vertebrates, including humans. In particular, human hepatitis B virus (HBV) is a leading cause of cirrhosis and hepatocellular carcinoma worldwide (3). In 2017, a new group of viruses distantly related to hepadnaviruses was identified in teleost fishes (4), and related viral sequences have since been reported (5). These viruses may represent a new viral family of pararetroviruses called the nackednaviruses, or *Nudnaviridae* (2). Their genome is slightly smaller than that of hepadnaviruses (2.7-3.1 kb versus 3.0-3.4 kb) and, importantly, they do not encode an envelope protein. The discovery of nackednaviruses suggested an ancient origin for hepadnaviruses from non-enveloped ancestors in fish (4).

In a recent study, we described DNA virus sequences from fecal samples of different bat species sampled in Spain (6). Analysis of the raw data from this study led us to identify a viral contig with an ambiguous taxonomic classification within the order Blubervirales. This sequence was obtained from the feces of a single *Myotis scalerai* individual captured on June 2022 in the Sima de la Higuera, a cave near the village of Pliego in the province of Murcia, Spain. PCR and sequencing of the cytochrome B gene confirmed that this sample was from *M. scalera*i. The viral contig corresponded to a complete circular genome of 3472 nt, as it had terminal redundancy, and was sequenced with an average coverage of 150 reads per base. Its identity was then confirmed by PCR using sequence-specific primers that allowed us to amplify the entire viral genome in overlapping fragments.

This virus, which we called Pliego virus, contained two ORFs of 1149 and 2376 nt homologous to, but highly divergent from, the capsid/core (C) and polymerase (P) of hepadnaviruses and nackednaviruses. BLASTp analysis indicated that the closest sequences for the C and P proteins corresponded to HBV (Genbank accession ANQ89943.1) and fish-associated hepatitis B virus (WAQ80622.1), respectively. However, the presence of multiple stop codons in the S-congruent reading frame revealed a typical nackednavirus feature. The Pliego virus genome contained three additional putative ORFs unrelated to those of other nackednaviruses or hepadnaviruses (**Figure 1A**), but the structure and function of the corresponding proteins could not be predicted using Phyre2 (7).

**Figure 1.**
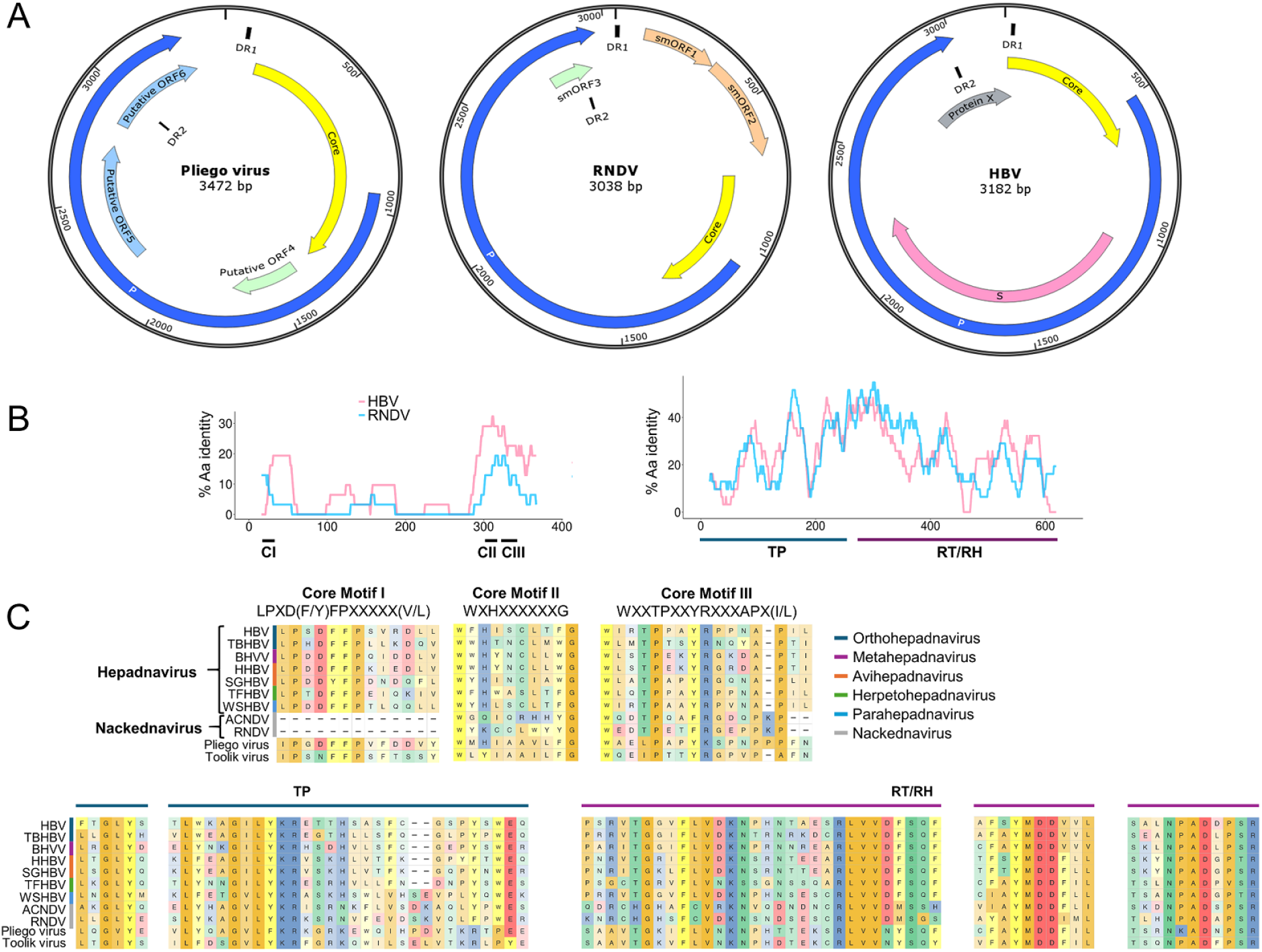
Comparative genome annotation. **A.** Genome organization of Pliego virus, RNDV, and HBV. In addition to genes with known function, putative ORFs and two short direct repeats of sequence AACTTCTACTGCAC (DR1 and DR2) are indicated. **B**. Similarity between the amino acid sequences of Pliego virus and RNDV or HBV for the C (left) and P (right) proteins, using a 30-residue sliding window. Core conserved motifs CI-CIII, terminal protein domain (TP), and reverse transcriptase and RNaseH domains (RT/RH) are shown. **C**. Alignments of the three core conserved motifs (mapping to residues 36-48, 303-312, and 326-341 of the C protein in the Pliego virus sequence) and selected regions of P domains for seven representatives of different hepadnavirus genera, two nackednaviruses, Pliego virus, and Toolik virus. Notice that, in the GLY conserved motif of TP, L is substituted for V in Pliego virus and I in Toolik virus. In contrast, the YMDD motif of the RT/RH is fully conserved.

A similarity plot of ORF C against a representative nackednavirus (rockfish nackednavirus, RNDV) and HBV showed a very high divergence (10.5% and 11.9% overall amino acid identity, respectively, **Figure 1B**). Nevertheless, we were able to identify conserved core motifs of vertebrate hepatitis B viruses (8), which were more similar to those of hepadnaviruses than to those of nackednaviruses (**Figure 1C**). The P protein was less extremely divergent than C (23.8% and 18.3% overall amino acid identity with RDNV and HBV, respectively; **Figure 1B**). In pararetroviruses, the P protein contains a reverse transcriptase (RT; IPR00477) and RNaseH (RH; IPR001462) domain, as well as a terminal protein (TP; IPR000201) domain involved in the initiation of reverse transcription (9), all of which were detected (**Figure 1C**). However, the RNA element epsilon, whose secondary structure is essential for the priming of reverse transcription (10), was not found. Similar to nackednaviruses (4), the spacer between the TP and RT found in hepadnaviruses was shorter than in hepadnaviruses. Other genomic motifs conserved in nackednaviruses and hepadnaviruses were identified, such as the direct repeats (DR1 and DR2) containing the TATA box and the polyadenylation signal.

We set out to detect similar sequences in databases. To this end, we first searched two large bat metagenomics projects (PRJNA953205, PRJNA929070), but this did not yield any hit. We then carried out a Blastp analysis of the C and P proteins in IMG/VR4, an extended database of uncultivated virus genomes (11). This revealed an unpublished 3316 nt viral sequence (Scaffold ID Ga0075039_1486360) assembled in a metagenomics study of permafrost soil carried out in the Toolik Field Station, Alaska. This “Toolik virus” shared amino acid sequence identities of 34.5% and 43.4% with Pliego virus C and P proteins, respectively, and also lacked a gene encoding an S protein. As for Pliego virus, we were not able to detect the RNA element epsilon (**Figure 1C**).

We then inferred the phylogenetic relationships between Pliego virus, Toolik virus hepadnaviruses, nackednaviruses, and other reverse-transcribing viruses (**Figure 2**). While nackednaviruses and hepadnaviruses showed well-supported distinct branches forming sister clades, a third cluster was formed by the Pliego and Toolik viruses, which occupied a basal position relative to all known animal pararetroviruses. Following previous work, we used the P protein sequence of an endogenous avihepadnaviral element (eAHBV-FRY) integrated into the Neoaves genome to estimate divergence times, assuming that the root of the eAHBV-FRY clade dated from the onset of the Neoaves radiation around 69-67 million years ago (4, 12, 13). The corresponding time-calibrated Bayesian tree indicated a divergence time of about 450 million years between nackednaviruses and hepadnaviruses, similar to the value obtained in previous work (4), while Pliego and Toolik viruses would have diverged more than 500 million years ago from the clade containing both nackednaviruses and hepadnaviruses. These results support the hypothesis that ancestral animal pararetroviruses were non-enveloped, and that the S ORF in hepadnaviruses probably arose by genetic overprinting (4).

**Figure 2.**
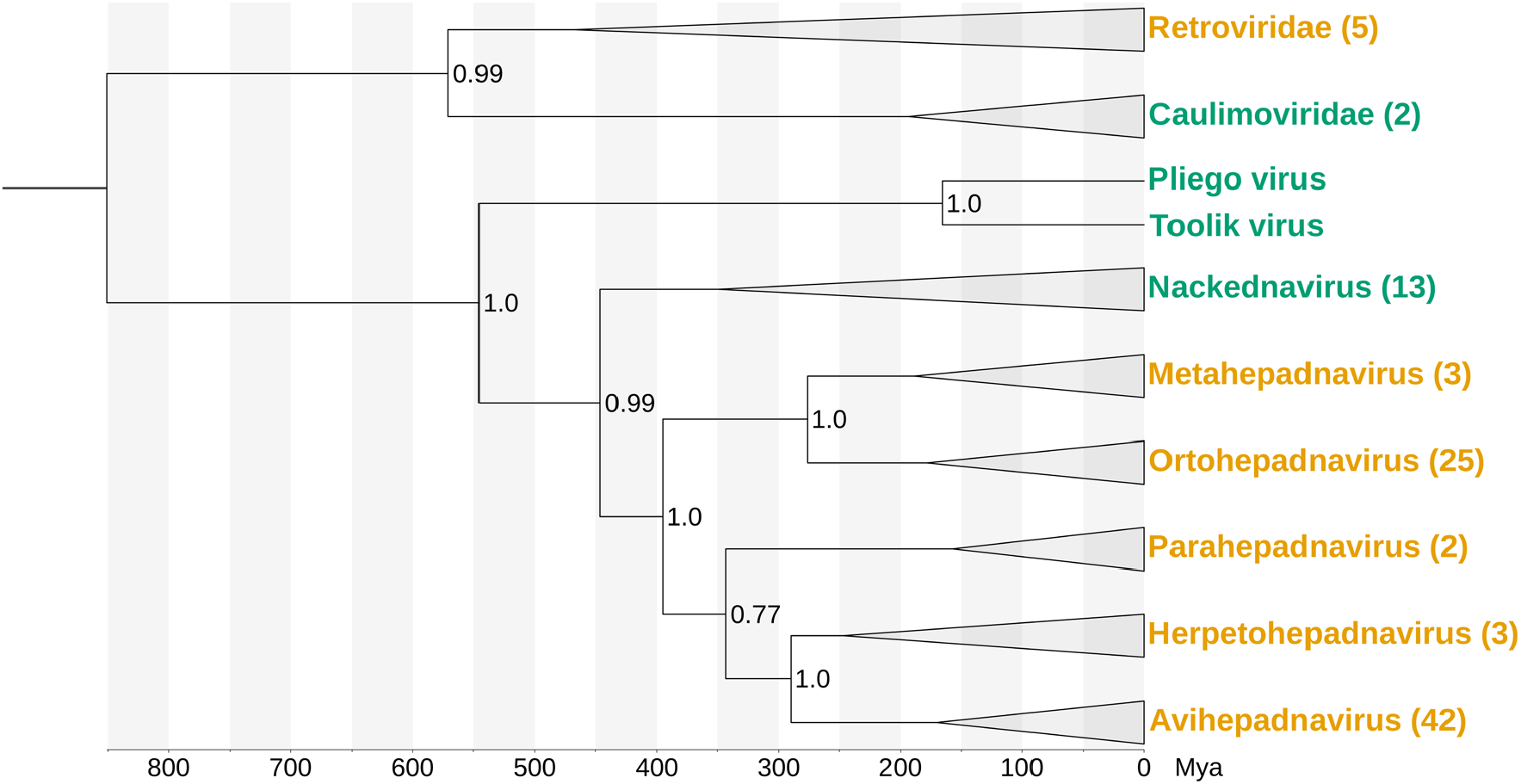
Phylogenetic position of the newly described cluster. Rooted Bayesian phylogenetic tree of the P protein. The Pliego virus and Toolik virus sequences were added to the alignment used in a previous work (4). Nodes are collapsed by genus or family, and the number of sequences in each clade is given in parentheses. Enveloped viruses are shown in yellow and non-enveloped viruses in green. Scale bar, millions of years ago (Mya). Numbers at branching nodes indicate posterior probability values.

Certain features are conserved between nackednaviruses and hepadnaviruses. For instance, both groups probably share a reverse transcription mechanism involving primer-independent initiation mediated by the epsilon RNA element (10). It was therefore surprising that we did not find this element in Pliego or Toolik virus. The element might be missing or too divergent to be detected. Another conserved feature is the capsid architecture (14), although nackednaviruses have T = 3 symmetry while hepadnaviruses have T = 4 symmetry. Since the genome sizes of Pliego and Toolik viruses are larger than those of the other nackednaviruses, we speculate that their structure may be more similar to that of hepadnaviruses, but structural analyses will be needed to clarify this.

Previous work has identified nackednaviruses exclusively in teleost fish (4, 5). In contrast, the Pliego virus sequence was obtained from the feces of a bat that feeds exclusively on arthropods (15), whereas the Toolik virus sequence was obtained from permafrost soil. Therefore, the newly identified viruses are unlikely to be restricted to fish. We can speculate that this new clade might be originally from arthropods and represent the ancestors of fish nackednaviruses, which subsequently evolved into vertebrate hepadnaviruses. However, cross-species transmission could alter this scheme. Although cross-species transmission is generally thought to be rare among DNA viruses (16, 17), it has been suggested for bat hepadnaviruses (18) and fish nackednaviruses (19). Alternatively, the Pliego virus might be a true bat virus, raising the question of whether it may pose a zoonotic risk, as bats are a known source of primate hepadnaviruses (20).

## Methods

### Sample collection and DNA extraction

Samples were obtained in a recent study as described previously (6). Briefly, individual bats were captured using nylon mist nets and a harp trap, identified visually to the species level, sexed, measured, weighed and briefly placed in cotton bags to recover fresh fecal samples. Samples were collected in tubes containing phosphate-buffered saline, kept cold initially, and then at -20 °C until they were transported to the laboratory and stored at -80 °C for further processing. Fecal samples were homogenized using ceramic beads and supernatants were filtered through a 1.2 µm pore size. Filtrates were used for total nucleic acid extraction and extracts were stored at -80 °C. For host taxonomic confirmation, we amplified by PCR and Sanger sequenced a 148 bp region of the cytochrome B gene using specific primers as described previously (21).

### Pliego virus sequencing

Sequencing libraries were prepared using the Nextera XT DNA kit and subjected to paired-end sequencing in a NextSeq 550 device with a 150 bp read length at each end. Reads were deduplicated, quality filtered with a quality trimming threshold of 20, and reads below 70 nucleotides in length were removed. De novo sequence assembly was performed using SPAdes v3.15.4 (22) with the meta option, and MEGAHIT v1.2.9 (23) using default parameters. Contig taxonomically classification was performed with Kaiju v1.9.0 (24). Virsorter2 v2.2.4 (25) was used to detect viral sequences, which were confirmed using CheckV v1.0.1 (26). Coverage statistics were calculated by remapping the trimmed and filtered reads from the sequencing library using Bowtie2 v2.2.5 (29). To confirm the sequence and completeness of Pliego virus genome, the following specific primers were designed to PCR-amplify six overlapping fragments of approximately 700 bp: 1F (5’-CGCGCTTATTCATGCTCA), 1R (5’-TTTGTGGTGGTGTTGGTGAT), 2F (5’-TCACACAAGGGTGGATACCAT), 2R (5’-TTCTTCGGGAGTTCCATACG), 3F (5’-CAAGAAAAATGGGAGGATGC) and 3R (5’-GACCGTCGGATATGGTGAGT), 4F (5’-TATCATCAGCCGCAGTAACG), 4R (5’-ACCCGATAAGTCGTTGATGG), 5F (5’-ACGAGACAAGGCAACCAAAT), 5R (5’-TTCGGTTGTGTCCATTTTCA), 6F (5’-CCCGTGCATTTCGTCTCTAT), and 6R (5’-CAGGCAATCAAGCGTTACAA).

### Sequence analysis and phylogenetic reconstruction

ORFs were predicted using ORFfinder (ncbi.nlm.gov/orffinder) and protein domains were annotated using InterProScan v5.63-95.0 (27) with the Pfam database v35.0. ORFs with missing protein domains were also analyzed with Phyre2 (7) to predict protein structure and function. Similarity plots for C and P protein alignments (sliding window of 31 residues) were created using R 4.4.0 and Biostrings v2.70.2 (https://bioconductor.org/packages/Biostrings). The Pliego virus and Toolik virus P sequences were aligned to those used in a previous study (4). The less conserved regions in the global alignment were removed using TrimAL v1.2rev59 (30) with the gappyout option, resulting in a final alignment of 490 positions. JTT+G4 was selected using ProtTest v3.4.2 (31) as the best-fitting amino acid substitution model. A time-calibrated Bayesian tree was computed with BEAST v2.7.6(28), using a calibrated Yules speciation prior and an optimized relaxed clock model with log-normal distribution. As previously described (4), a uniform mean clock rate and a normally distributed prior with mean of 69.2 and standard deviation of 1.735 for the age of the eAHBV-FRY root were used. The final time-calibrated tree was computed combining three independent analyses using a chain length of 50 million each.

## Ethics statement

Bats were captured in accordance with the European directive regulating the protection of animals used for scientific research (2010/62/EU, Article 1), subsequently transposed into Spanish legislation (Royal decree 53/213, 1 February, Article 2). The procedures followed in this study (i.e. capture, non-invasive handling and in situ release of wild animals) are not under the status of animal experimentation and hence do not require an IACUC approval document, but instead specifically a permit from the competent regional authority (Ref. Exp. 2022-VS (FAU22_009)).

## Acknowledgments

This research was financially supported by grant PID2020-118602RB-I00 from the Spanish Ministerio de Ciencia e Innovación (MICINN) to R.S and J.-M.C., grant CIAICO/2022/110 from the Conselleria de Educación, Universidades y Empleo (Generalitat Valenciana) to R.S., and ERC Advanced Grant 101019724-EVADER to R.S.

## Data availability

The raw sequence reads were deposited in the Sequence Read Archive of GenBank under accession numbers SRR27912327-51. The new viral contig described in this study was deposited in Genbank under accession number PQ119727.

## References

1. P.-Y. Teycheney, et al., ICTV Virus Taxonomy Profile: Caulimoviridae. J. Gen. Virol. 101, 1025–1026 (2020).

2. L. Magnius, et al., ICTV Virus Taxonomy Profile: Hepadnaviridae. J. Gen. Virol. 101, 571–572 (2020).

3. W.-J. Jeng, G. V. Papatheodoridis, A. S. F. Lok, Hepatitis B. Lancet Lond. Engl. 401, 1039–1052 (2023).

4. C. Lauber, et al., Deciphering the Origin and Evolution of Hepatitis B Viruses by Means of a Family of Non-enveloped Fish Viruses. Cell Host Microbe 22, 387-399.e6 (2017).

5. C. E. Ford, C. D. Dunn, E. M. Leis, W. A. Thiel, T. L. Goldberg, Five Species of Wild Freshwater Sport Fish in Wisconsin, USA, Reveal Highly Diverse Viromes. Pathog. Basel Switz. 13, 150 (2024).

6. J. Buigues, et al., Full-genome sequencing of dozens of new DNA viruses found in Spanish bat feces. Microbiol. Spectr. e0067524 (2024). 10.1128/spectrum.00675-24.

7. L. A. Kelley, S. Mezulis, C. M. Yates, M. N. Wass, M. J. E. Sternberg, The Phyre2 web portal for protein modeling, prediction and analysis. Nat. Protoc. 10, 845–858 (2015).

8. J. A. Dill, et al., Distinct Viral Lineages from Fish and Amphibians Reveal the Complex Evolutionary History of Hepadnaviruses. J. Virol. 90, 7920–7933 (2016).

9. D. N. Clark, J. M. Flanagan, J. Hu, Mapping of Functional Subdomains in the Terminal Protein Domain of Hepatitis B Virus Polymerase. J. Virol. 91, e01785–16 (2017).

10. J. Beck, S. Seitz, C. Lauber, M. Nassal, Conservation of the HBV RNA element epsilon in nackednaviruses reveals ancient origin of protein-primed reverse transcription. Proc. Natl. Acad. Sci. U. S. A. 118, e2022373118 (2021).

11. A. P. Camargo, et al., IMG/VR v4: an expanded database of uncultivated virus genomes within a framework of extensive functional, taxonomic, and ecological metadata. Nucleic Acids Res. 51, D733–D743 (2023).

12. A. Suh, J. Brosius, J. Schmitz, J. O. Kriegs, The genome of a Mesozoic paleovirus reveals the evolution of hepatitis B viruses. Nat. Commun. 4, 1791 (2013).

13. S. Claramunt, J. Cracraft, A new time tree reveals Earth history’s imprint on the evolution of modern birds. Sci. Adv. 1, e1501005 (2015).

14. S. Pfister, et al., Structural conservation of HBV-like capsid proteins over hundreds of millions of years despite the shift from non-enveloped to enveloped life-style. Nat. Commun. 14, 1574 (2023).

15. R. Novella-Fernandez, et al., Trophic resource partitioning drives fine-scale coexistence in cryptic bat species. Ecol. Evol. 10, 14122–14136 (2020).

16. J. L. Geoghegan, S. Duchêne, E. C. Holmes, Comparative analysis estimates the relative frequencies of co-divergence and cross-species transmission within viral families. PLoS Pathog. 13, e1006215 (2017).

17. V. A. Costa, et al., Limited cross-species virus transmission in a spatially restricted coral reef fish community. Virus Evol. 9, vead011 (2023).

18. F.-Y. Nie, et al., Extensive diversity and evolution of hepadnaviruses in bats in China. Virology 514, 88–97 (2018).

19. V. A. Costa, et al., Host adaptive radiation is associated with rapid virus diversification and cross-species transmission in African cichlid fishes. Curr. Biol. CB 34, 1247-1257.e3 (2024).

20. J. F. Drexler, et al., Bats carry pathogenic hepadnaviruses antigenically related to hepatitis B virus and capable of infecting human hepatocytes. Proc. Natl. Acad. Sci. U. S. A. 110, 16151–16156 (2013).

21. A. Lopez-Oceja, D. Gamarra, S. Borragan, S. Jiménez-Moreno, M. M. de Pancorbo, New cyt b gene universal primer set for forensic analysis. Forensic Sci. Int. Genet. 23, 159–165 (2016).

22. S. Nurk, D. Meleshko, A. Korobeynikov, P. A. Pevzner, metaSPAdes: a new versatile metagenomic assembler. Genome Res. 27, 824–834 (2017).

23. D. Li, C.-M. Liu, R. Luo, K. Sadakane, T.-W. Lam, MEGAHIT: an ultra-fast single-node solution for large and complex metagenomics assembly via succinct de Bruijn graph. Bioinforma. Oxf. Engl. 31, 1674–1676 (2015).

24. P. Menzel, K. L. Ng, A. Krogh, Fast and sensitive taxonomic classification for metagenomics with Kaiju. Nat. Commun. 7, 11257 (2016).

25. J. Guo, et al., VirSorter2: a multi-classifier, expert-guided approach to detect diverse DNA and RNA viruses. Microbiome 9, 37 (2021).

26. S. Nayfach, et al., CheckV assesses the quality and completeness of metagenome-assembled viral genomes. Nat. Biotechnol. 39, 578–585 (2021).

27. P. Jones, et al., InterProScan 5: genome-scale protein function classification. Bioinforma. Oxf. Engl. 30, 1236–1240 (2014).

28. R. Bouckaert, et al., BEAST 2.5: An advanced software platform for Bayesian evolutionary analysis. PLoS Comput. Biol. 15, e1006650 (2019).

29. B. Langmead, S. L. Salzberg, Fast gapped-read alignment with Bowtie 2. Nat Methods 9, 357–359 (2012).

30. S. Capella-Gutiérrez, J. M. Silla-Martínez, T. Gabaldón, trimAl: a tool for automated alignment trimming in large-scale phylogenetic analyses. Bioinforma. Oxf. Engl. 25, 1972–1973 (2009).

31. D. Darriba, G. L. Taboada, R. Doallo, D. Posada, ProtTest 3: fast selection of best-fit models of protein evolution. Bioinforma. Oxf. Engl. 27, 1164–1165 (2011).

